# Evolution of the ribbon-like organization of the Golgi apparatus in animal cells

**DOI:** 10.1101/2023.02.16.528797

**Authors:** Giovanna Benvenuto, Serena Leone, Emanuele Astoricchio, Sophia Bormke, Sanja Jasek, Enrico D’Aniello, Maike Kittelmann, Kent McDonald, Volker Hartenstein, Valentina Baena, Héctor Escrivà, Stephanie Bertrand, Bernd Schierwater, Pawel Burkhardt, Iñaki Ruiz-Trillo, Gáspár Jékely, Jack Ullrich-Lüter, Carsten Lüter, Salvatore D’Aniello, Maria Ina Arnone, Francesco Ferraro

**Author notes:** These authors contributed equally (Giovanna Benvenuto, Serena Leone).

## Abstract

The Golgi ribbon is a structural organization formed by linked Golgi stacks that is believed to be exclusive to vertebrate cells. Its functional contribution to cellular processes is unclear, yet its disruption is associated with several human pathologies. In this study we address the evolutionary origin of the Golgi ribbon, describe a potential molecular mechanism for its emergence and identify a cellular process in which it may be involved. We observed the ribbon-like architecture in the cells of several metazoan taxa, suggesting its early appearance during animal evolution before the emergence of vertebrates. Supported by AlphaFold2 modelling, we propose that the evolution of the complex between two Golgi resident proteins, Golgin-45 and GRASP, led to the tethering of Golgi stacks into the ribbon-like configuration. Finally, we find that the ribbon is assembled during the early embryogenesis of deuterostome animals, a strong indication of its role in development. Overall, our study indicates that the Golgi ribbon is functionally relevant beyond vertebrates and calls for further investigations to decipher its elusive functions.

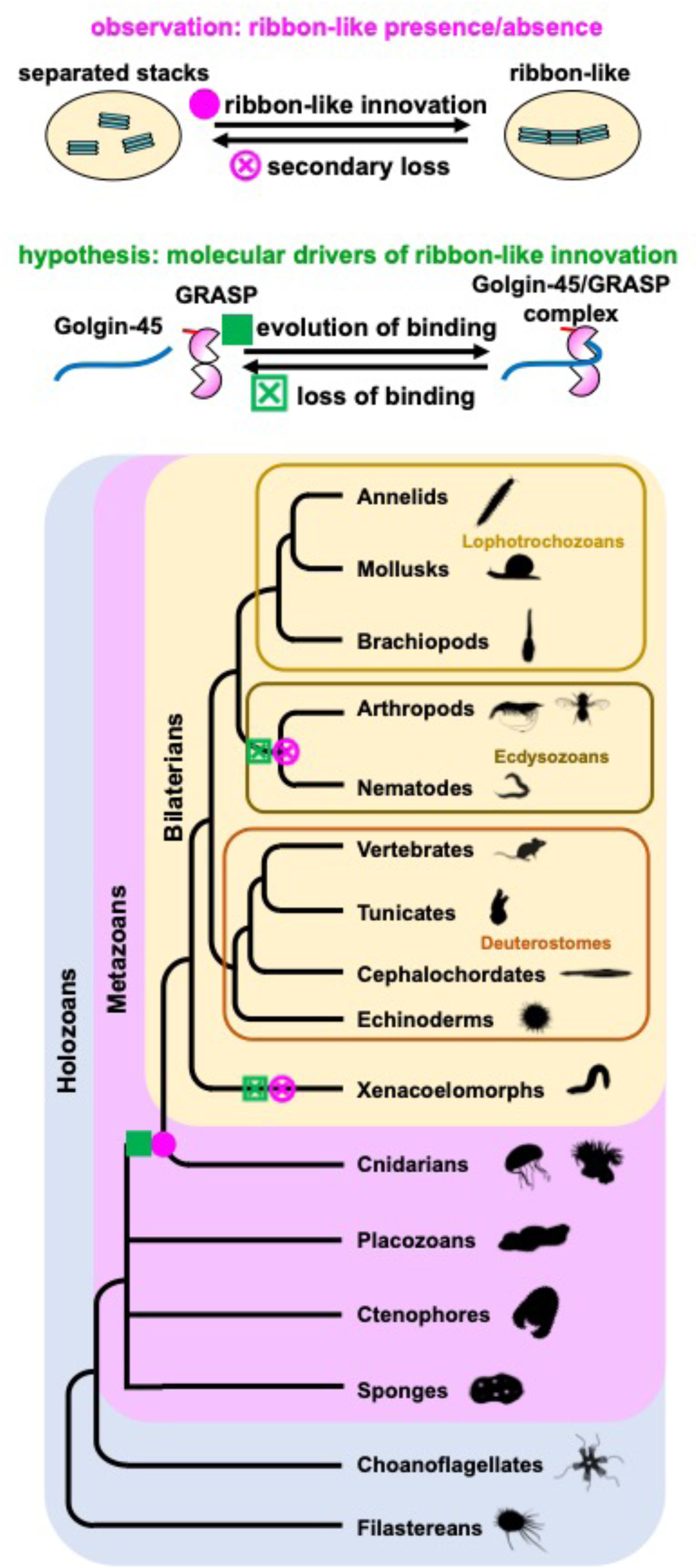
Graphical abstract.

## Introduction

Sitting at the center of the exocytic pathway, the Golgi apparatus is involved in the processing and sorting of secretory cargoes. This biosynthetic function remains the most actively investigated^1–3^, but recent evidence indicate that the Golgi is also involved in secretion-independent cellular processes, such as stress sensing and signaling, apoptosis, autophagy, proteostasis and innate immunity^4–11^. The Golgi’s fundamental structural unit is the stack, formed by a pile of flat membrane saccules, known as cisternae. Across eukaryotes, the Golgi occurs as a single- or multi-copy organelle, depending on the number of stacks per cell. In animals, when present as multi-copy organelle, the Golgi is observed in two configurations: stacks either remain separate or link to each other into a single centralized structure that was first described as “ribbon-like” by Camillo Golgi^12^. The current consensus is that the Golgi ribbon is present only in vertebrate cells. Despite having been widely investigated, for the most part in cultured mammalian cells, the biological functions of the Golgi ribbon still remain obscure^13–15^ and it is unclear which selective pressures might have led to the evolution of this Golgi configuration. During mitosis, mammalian cells disassemble and reassemble the ribbon in a finely tuned process^16^; such a level of regulation suggests that the ribbon architecture must be functionally important. This conclusion is supported by the existence of several pathologies in which ribbon breakdown (Golgi “fragmentation”) is a morphological phenotype, most notably neurodegenerative diseases but also cancer and viral infections^17–23^. Therefore, deciphering the roles that the Golgi ribbon plays in cellular physiology may also help us understand the pathological consequences stemming from its disruption. In this study, through a comparative approach borrowed from evolutionary studies, we asked three questions regarding the ribbon architecture of the Golgi apparatus. 1) When did it appear during animal evolution? 2) Which molecules might have mediated this structural innovation? 3) Which biological functions does it carry out? We answer the first one; propose a testable hypothesis for the second one; and, regarding the third one, we produce experimental data that point toward development as a biological process in which the ribbon may play a functional role.

## Results

### A ribbon-like Golgi complex is common in animals

According to the current consensus, only vertebrate cells form Golgi ribbons. Therefore, we were intrigued by morphological data showing Golgi centralization in the embryos of two sea urchin species, *Strongylocentrotus purpuratus* and *Lytechinus variegatus*^24,25^, hinting at the possibility that the ribbon organization of the Golgi might be more common than thought at present. To assess if this is the case, we surveyed the Golgi ultrastructure in representatives of several animal taxa and closely related unicellular eukaryotes, which, together, comprise the eukaryotic holozoan clade. We had to distinguish between *bona fide* ribbon and ribbon-like architectures. In mammalian cells the Golgi ribbon is characterized by some level of membrane continuity between cisternae of adjacent stacks, which can be detected by fluorescent recovery after photobleaching assays and electron tomography^26,27^. As our survey relied for the most part on electron micrographs of thin sections where membrane continuity between cisternae of juxtaposed stacks cannot be easily assessed, we describe Golgi organizations reminiscent of the mammalian ribbon as ribbon-like. Golgi stack dimers have been observed in *Drosophila melanogaster* cells^28^, which have dispersed Golgi elements^29^, and even in mammalian cells after ribbon unlinking by microtubule depolymerization^30^. Therefore, we identified Golgi complexes as “ribbon-like” only when three or more closely apposed stacks were observed in electron micrographs (Figure 1A). In mammalian cell types, the ribbon architecture, though common, is not ubiquitous. Differentiated tissues such as muscles, acid-secreting gastric cells, and spinal ganglion neurons, for instance, display Golgi complexes made by separated stacks^31–33^. For this reason, wherever possible, several cell types of the organisms under consideration were inspected and, whenever available, we supplemented our survey with morphological evidence from the literature.

**Figure 1.**
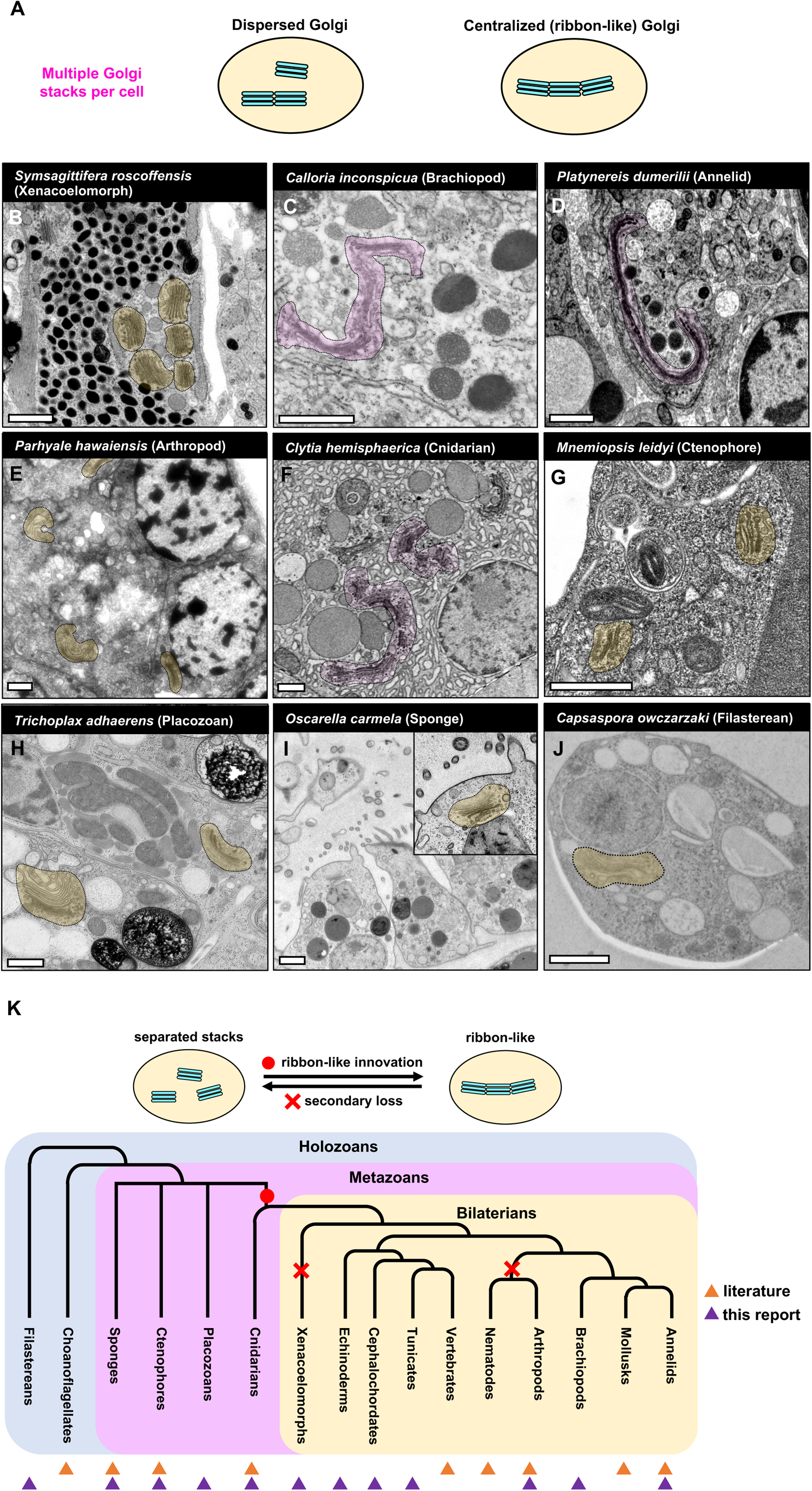
Golgi architecture in holozoans. The Golgi organization in holozoan exemplars from diverse taxa was analyzed at the ultrastructural level; separated and linked stacks are highlighted in light ochre and light magenta, respectively. (A) Observation of three or more Golgi stacks in close contact and/or with membrane continuities is the criterion adopted for positive identification of the ribbon-like configuration in electron micrographs. (B) The xenacoelomorph *Symsagittifera roscoffensis* (Roscoff worm), secretory cell. (C) The brachiopod *Calloria inconspicua*, epidermal cell of the mantle lobe of the three-lobed larva. (D) The annelid *Platynereis dumerilii*, glial cell of the 3-day old larva. (E) The crustacean *Parhyale hawaiensis*, nerve cell. (F) The cnidarian *Clytia hemisphaerica*, gonad gastrodermal cells. (G) The ctenophore *Mnemiopsis leydi*, epithelial cells. (H) The placozoan *Tricoplax adhaerens*. (I) The sea sponge *Oscarella carmela*, choanocyte. (J) The filasterean *Capsaspora owczarzaki.* Scale bars: 1 μm. (K) Deduced evolutionary emergence of the ribbon-like Golgi organization. Ribbon-like absence in both arthropods and nematodes, which both belong to the ecdysozoan superphylum, may indicate that loss of Golgi stack linking occurred in their common ancestor. See also Figure S1 and Movie S1.

We first looked at bilaterians. In the cells of the marine worm *Symsagittifera roscoffensis* (xenacoelomorph), separated stacks were observed (not shown). Interestingly, some of its secretory cells displayed closely apposed, though clearly distinct, Golgi stacks: an intermediate organization between separated Golgi elements and a ribbon-like organization (Figures 1B and S1A). Ribbon-like Golgis were found in epidermal cells of the three-lobed larva of the brachiopod *Calloria inconspicua* and in several cell types of the marine annelid *Platynereis dumerilii* (Figures 1C, 1D, S1B and Movie S1). As Ramón y Cajal described ribbon-like Golgis in neurons and epithelial cells of the common earthworm^34^, we conclude that this Golgi organization is common among annelids. In mollusks, a centralized Golgi that fragments at mitosis was observed in spermatocytes of the snail *Paludina vivipara* more than a century ago^35^, while other reports show ribbon-like Golgi complexes in other species (e.g., *Helix pomatia*^36^ and *Helix aspersa*^37^). Cells of the roundworm *Caenorhabditis elegans*, a nematode, and of the fruit fly *Drosophila melanogaster*, an arthropod, two model organisms widely used in genetics and cell biology, display Golgi complexes consisting of several, separated stacks^28,29,38,39^. To test whether Golgi stack decentralization is an arthropod feature, as opposed to *Drosophila*/insect-specific, we analyzed the ultrastructure of the crustacean *Parhyale hawaiensis*, observing separated stacks in neurons (Figure 1E) and in all other inspected cell types (not shown). It is therefore likely that a decentralized Golgi is the typical configuration in arthropods, not just of *Drosophila* and other insects (e.g., bees, aphids, and mosquitos^40–42^). We then analyzed cnidarians: in the hydrozoan *Clytia hemisphaerica*, the secretory gland cells of the gastroderm, but not other cell types, display stacks linked into a ribbon-like structure (Figure 1F), which is also observed in phagocytic cells of another cnidarian, the actinia *Phelliactis robusta*^43^. In the ctenophore *Mnemiopsis leidyi*, epithelial and comb cells (Figures 1G and S1C), nerve net neurons, mesogleal neurons, and sensory cells^44,45^ all display separated stacks. Among other animals, we found a single Golgi stack in all cells of two placozoan species: *Trichoplax adhaerens* (Figures 1H, S1D and S1E) and *Hoilungia hongkongensis*^46^. Like placozoans, the sea sponge *Oscarella carmela* (Figure 1I and reference^47^) and other species (genera *Chondrosia*, *Crambe* and *Petrosia*; not shown) have a single Golgi stack per cell. In choanoflagellates and filastereans, which are unicellular holozoans, the Golgi is also present as a single stack per cell (Figure 1J and references^47,48^).

Vertebrates (chordates) and echinoderms (ambulacraria) belong to the deuterostome clade of bilaterian animals. We assessed whether the Golgi ribbon is found in non-vertebrate deuterostomes, investigating the ultrastructure of a sea urchin, a tunicate and a cephalochordate species. Indeed, we find that all three species display ribbon-like Golgis (see section “Developmental assembly of the Golgi ribbon”, below).

In summary, despite a relatively small sampling (Figure S1F), ribbon-like Golgi complexes are easily observed in cells of cnidarians and bilaterians, and not found outside these animal taxa. Obviously, the presence of multiple stacks per cell is a precondition for their clustering and ribbon formation, but it appears to be not sufficient. Point in case are sponges, which usually display a single Golgi stack per cell (Figure 1I and reference^47^); however, in rare instances, such as the gemmule’s spongocytes of the freshwater species *Ephydatia fluviatilis*^49^, sponge cells with multiple stacks are observed, but remain separate. It should be noted that in our survey, except for *Platynereis dumerilii* and *Strongylocentrotus purpuratus* (Figure S1B, Movie S1 and Figure S4E) for which volume electron microscopy was available, the thin section electron micrographs did not allow to assess whether all the Golgi stacks in a cell are linked into a single ribbon-like organization or form multiple “mini-ribbons”. Nonetheless, in those cases where we identified a ribbon-like organization we can state that the process of stack centralization is clearly observed. In conclusion, the most parsimonious explanation accounting for our results and the literature’s data is that the ribbon-like Golgi likely evolved in the common ancestor of cnidarians and bilaterians, and was secondarily lost in xenacoelomorphs, arthropods, and nematodes (Figure 1K).

### Putative molecular mediators of ribbon-like Golgi emergence

Next, we asked which molecular innovations might have driven the emergence of the ribbon-like Golgi organization. If, as our survey suggests, this was a single evolutionary event, conservation of the molecular mechanisms of its formation would be expected. Among the several factors involved in the formation of the mammalian Golgi ribbon^50^, the molecular tethers GRASP (Golgi Reassembly and Stacking Protein) and the coiled-coil proteins collectively known as Golgins play a central role (Figure 2A)^50–56^. GRASPs comprise an evolutionarily conserved GRASP region, made of two atypical PDZ domains in tandem, and a more variable C-terminal unstructured region (Figures S2A and S2B). While GRASPs are encoded by a single gene in most eukaryotes, a duplication gave rise to two paralogs in jawed vertebrates (Figure S2C Data S1). Involved in several cellular processes^57–60^, GRASPs are capable of self-interaction and while they were initially considered to promote cisternal adhesion within the stack^61–63^, recent work unequivocally showed that they mediate Golgi stack tethering and ribbon formation but not cisternal stacking^55,56^. The two mammalian paralogs, GRASP55 and GRASP65, are recruited to Golgi membranes by myristoylation of the glycine in position 2^61,64^, which is conserved across holozoans (Data S1), and by interaction with Golgin-45 and GM130, respectively (Figure 2A)^65,66^. Such dual-anchoring is required for ribbon formation as it spatially orients GRASPs and allows their homo-dimerization/oligomerization in *trans*, thus tethering membranes of distinct Golgi stacks and promoting ribbon formation^67–69^. Golgins mediate vesicular traffic specificity^70–72^ and their knockdown results in secretory defects and ribbon unlinking into constituent stacks^50,53^; though, it is not clear whether these two phenotypes depend on the same or separate functions that Golgins play. The *Golgin-45* gene is an innovation of holozoan eukaryotes^73^. In mammals, the Golgin-45 protein interacts with the GRASP paralog GRASP55 (Figure 2A)^66^; and in cultured cells, either Golgin-45 knockdown or long-term degron-induced ablation of GRASP55, but not of GRASP65, result in Golgi ribbon unlinking^56,66^. As GRASP55 is more similar to the single GRASP proteins present in non-vertebrate bilaterians and cnidarians (Figure S2C), it is plausible that evolution of GRASP binding by Golgin-45 may have led to GRASP-mediated stack tethering (i.e., centralization) and ribbon-like Golgi evolution (Figure 2B). We searched and identified the Golgin-45 homologs in representatives of metazoans and of their closer unicellular relatives, confirming previous findings that this protein evolved in the common ancestor of holozoans^73^. In addition, we found that the *Golgin-45* gene was lost in choanoflagellates, and, among metazoans, in most xenacoelomorphs (Data S2), which interestingly do not display ribbon-like Golgi (Figures 1B and S1A).

**Figure 2.**
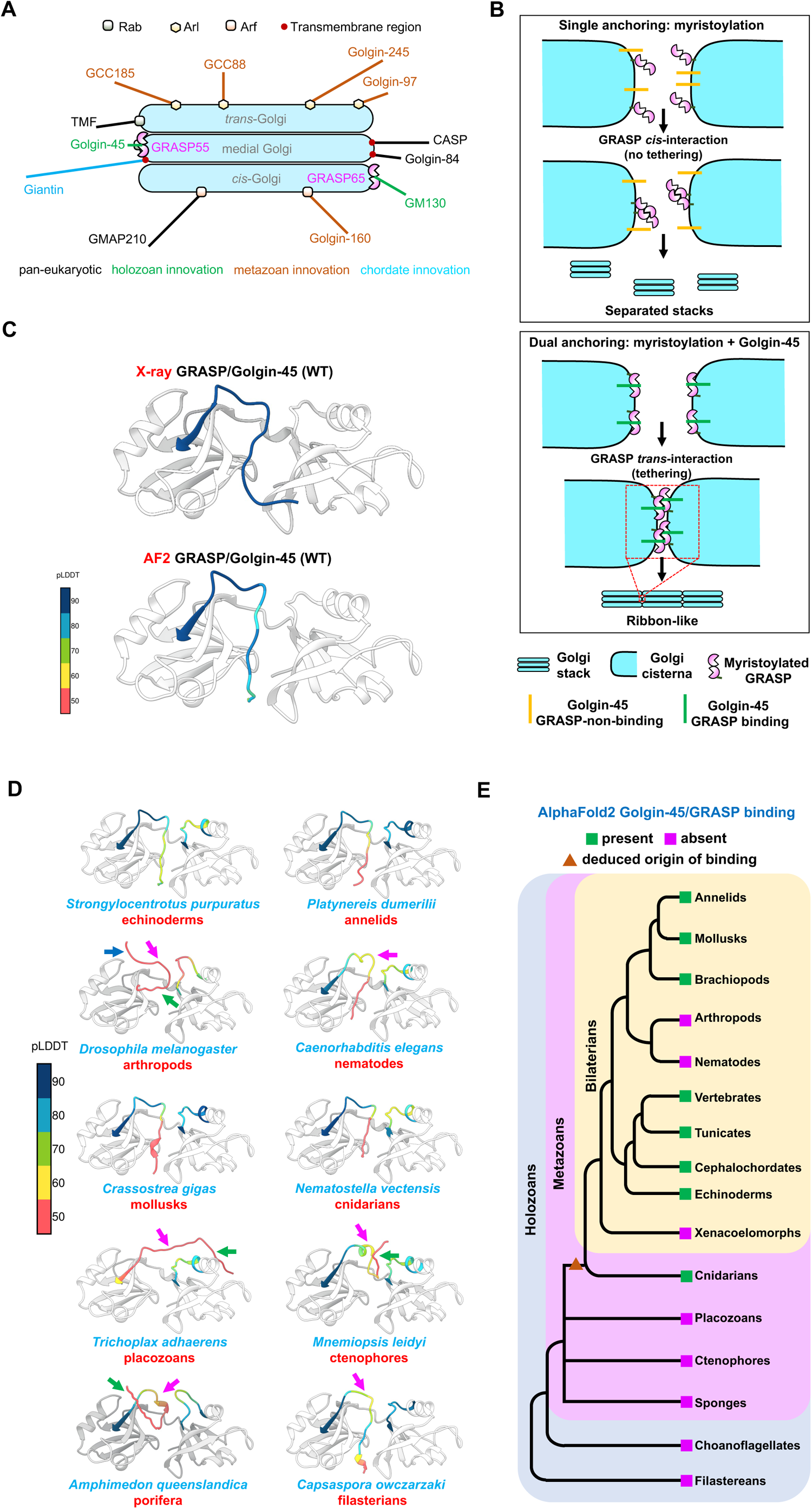
Putative molecular mediators of the ribbon-like Golgi emergence. (A) The Golgi localized molecular tethers Golgins and GRASPs. Golgins are coiled-coil proteins that localize to Golgi membranes by a transmembrane region or through recruitment by small GTPases of the Rab, Arl and Arf families. Golgin localization within the stack (see references^50,52,53,94,95^), their sizes (human homologs, bar length), and their evolutionary emergence^73^ are indicated. (B) Evolution of GRASP-mediated Golgi stack tethering. In mammalian cells, dual anchoring of GRASPs on Golgi membranes is required for self-interaction in *trans* and stack tethering. GRASP myristoylation is ancestral (see text and Data S1) and therefore its dual anchoring to Golgi membrane should originate from the evolution of binding by Golgin-45, leading to the emergence of stack linking and ribbon formation. (C) Solved structure (X-ray; PDB accession code 5H3J) and the AlphaFold2 (AF2) model of the mouse Golgin-45/GRASP complex. AF2 predicted structure almost overlaps the experimentally solved one (RMSD 3.040 Å for the Cα of the last 16 amino acids of the Golgin-45 C-terminal peptide). (D) AF2 models of holozoan GRASPs in complex with their conspecific Golgin-45 C-termini. Altered conformations, with respect to the mouse complex, are indicated by the arrows (blue for the PDZ-binding motif, red for the cysteine pair and green for the GRASP groove-interacting residues). (E) Presence and inferred evolutionary origin of GRASP binding by the C-terminus of Golgin-45, as deduced by AlphaFold2 modelling of complexes. See also Figures S2 and S3, Data S1 and S2.

The crystal structure of the complex between the C-terminus of mouse Golgin-45 and the GRASP domain of the conspecific GRASP55 has been solved^74^, highlighting the existence of three main interaction sites between the two proteins: i) a PDZ-binding motif spanning the four C-terminal amino acids of Golgin-45; ii) an atypical Zinc finger composed by two cysteines of Golgin-45 and a cysteine and a histidine in the GRASP domain; iii) the insertion of nine residues of Golgin-45 into the hydrophobic groove between the two PDZ domains of GRASP55 (Figure S2D)^74^. Binding experiments showed that the PDZ-binding motif and the cysteine pair are necessary for Golgin-45/GRASP complex formation, whereas the contribution of the groove-interacting residues remains unclear^74^ (Supplemental Results and Discussion). We aligned the C-terminal sequences of holozoan Golgin-45 proteins to assess conservation of the amino acids involved in GRASP interaction (Figure S2E). The PDZ binding motif and the cysteine pair are highly conserved, with the notable exception of *Drosophila melanogaster* and *Parhyale hawaiensis*, whose cells lack ribbon-like Golgi organization (Figure S2E, refs.^28,29^ and Figure 1E), whereas the Golgin-45 residues corresponding to those that interact with the GRASP groove are more variable across holozoans (Figure S2E).

As the untemplated AlphaFold2 model^75,76^ of the mouse Golgin-45/GRASP complex was generated with high confidence and was very similar to the crystal structure (Figure 2C and Supplemental Results and Discussion), we reasoned that Golgin-45/GRASP interactions may be predicted^77,78^ by modelling complexes of conspecific protein pairs. We considered binding to occur when GRASP interaction with the PDZ-binding motif and formation of the Zinc finger could be detected in the modelled complex. In bilaterians and cnidarians, all models predicted binding, except for arthropods (*Drosophila melanogaster* and *Parhyale hawaiensis*), nematodes (*Caenorhabditis elegans*) and the only xenacoelomorph species in which the *Golgin-45* gene is still present, *Hofstenia miamia* (Figures 2D and S3A). Golgin-45 was not predicted to bind its conspecific GRASP in ctenophores, porifera and unicellular filastereans (Figure 2D). The reliability of AlphaFold2 predictions was corroborated by modeling the complexes of various point mutants of the mouse Golgin-45 PDZ-binding motif, cysteine pair, and groove interacting residues that had been experimentally tested in *in vitro* GRASP binding assays^74^. The models obtained were consistent with the experimental data by those authors (Figure S3B and Supplemental Results and Discussion). We therefore deduce that a stable Golgin-45/GRASP interaction is present in cnidarians and bilaterians, likely appearing in their common ancestor, but was impaired by subsequent amino acid mutations in the Golgin-45 proteins of arthropods, nematodes, and in the only extant one of xenacoelomorphs (Figure S3A). In conclusion, AlphaFold2 models lend support to our hypothesis that the evolution of GRASP binding by Golgin-45 may have driven the appearance of stack tethering and the emergence of the ribbon-like Golgi organization (Figure 2E).

### Developmental assembly of the Golgi ribbon

As published morphological data were indicative of Golgi centralization in sea urchins^24,25^, we analyzed Golgi dynamics in the embryo of the sea urchin species *Paracentrotus lividus*. Time-course analysis of a fluorescent Golgi reporter showed that early in development, throughout the cleavage stage, the Golgi is present as separate elements which then cluster into centralized structures before hatching of the blastula (Figures 3A, S4A and S4B). Golgi clustering is rapid: within one hour, Golgi elements increase 10-fold in size while their number decreases 3-fold (Figures 3B, 3C and Movie S2). Afterwards, centralized Golgi complexes are observed in all cells of the embryo and at all developmental stages up to the planktonic pluteus larva (Figures 3A and S4C) and confocal imaging at higher magnification of post-clustering stages showed a morphology strongly reminiscent of the Golgi ribbon as observed in mammalian cells (Figure S4D). At the ultrastructural level, the arrangement of the sea urchin Golgi elements recapitulated confocal microscopy observations. Separated stacks cluster and finally establish connections with each other during early development, confirming that sea urchins centralize their Golgi apparatus into a ribbon-like architecture (Figure 3D). Centralized Golgi complexes were previously observed in the gastrula and prism stages of *Strongylocentrotus purpuratus*^24^. Indeed, we also observed ribbon-like Golgi complexes in the pluteus larva of this sea urchin species (Figure S4E). Like the ribbon of mammalian cells, sea urchin’s centralized Golgi undergoes disassembly/reassembly cycles during mitosis (Figure S4F) and its maintenance requires an intact microtubule network (Figures S4G and S4H)^14,16, 79–81^. All these characteristics strongly indicate that the centralized Golgi complexes of sea urchin cells are *bona fide* ribbons. As echinoderm representatives, sea urchins form part of the sister group to all chordates, including vertebrates. Therefore, the mechanisms mediating formation of the Golgi ribbon and its developmental timing might be conserved across the deuterostome clade. Indeed, we observed developmental Golgi stack clustering and ribbon-like formation in the cells of two non-vertebrate chordates, the sea squirt *Ciona robusta* (tunicate) and the lancelet *Branchiostoma lanceolatum* (cephalochordate) (Figures 3E and 3F). Golgi centralization also occurs in mouse embryos during the preimplantation stage^82^, indicating that the developmental ribbon assembly is indeed a conserved feature across deuterostomes. Such a Golgi dynamics suggests that the developmental switch from separated Golgi stacks to the ribbon-like configuration might play an important role during the initial stages of embryogenesis.

**Figure 3.**
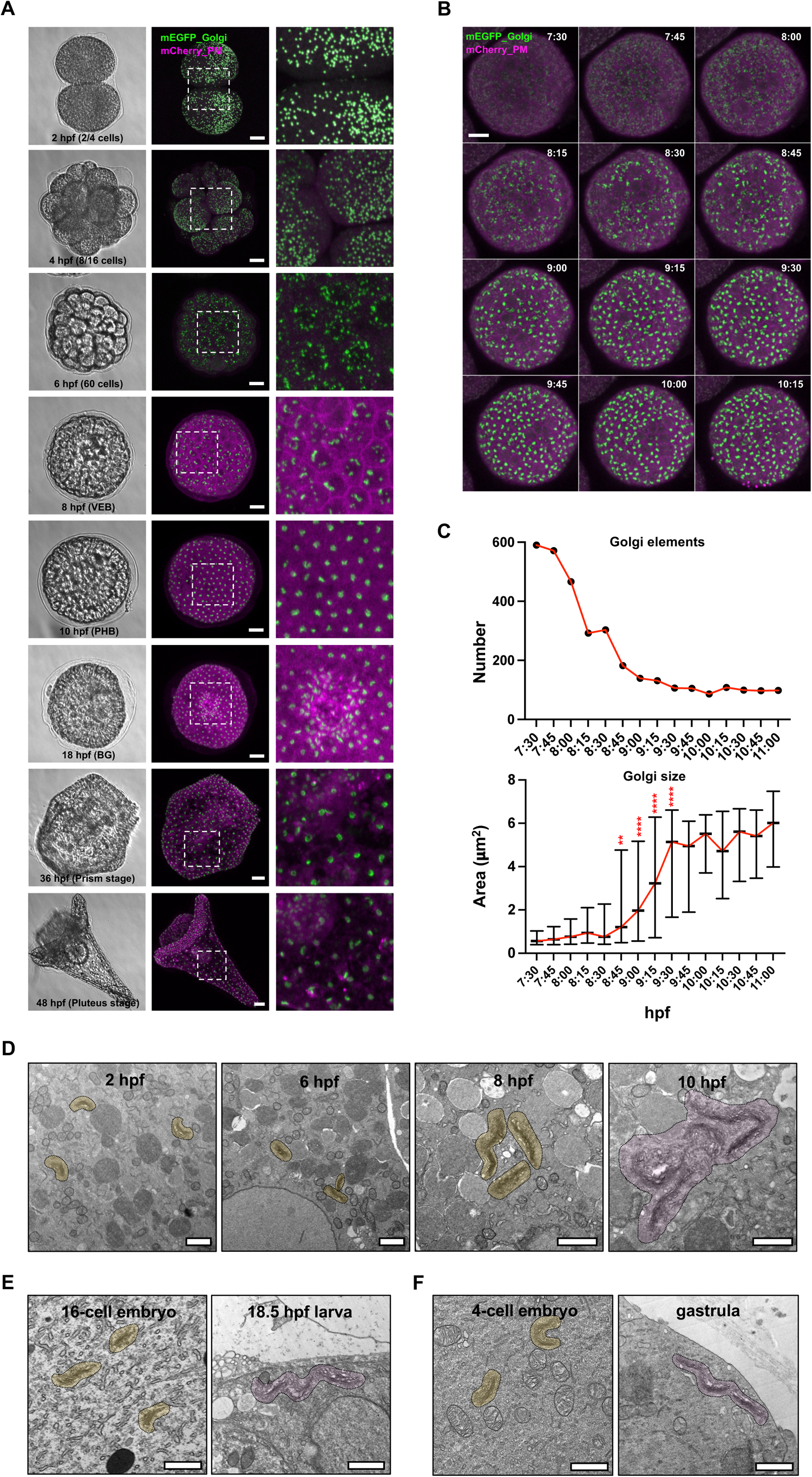
Developmental assembly of the Golgi ribbon. (A) Embryos of the sea urchin *Paracentrotus lividus* expressing fluorescent reporters of the Golgi apparatus and the plasma membrane (PM) were imaged at the indicated stages (hpf, hours post-fertilization; VEB, very early blastula; PHB, post-hatching blastula; BG, blastopore gastrula) by bright field and confocal microscopy (maximum intensity projections); right panels show magnifications of the middle panel insets; scale bars: 20 μm. (B) Maximum intensity projections of time-lapse confocal microscopy of an embryo microinjected as described in (A) and imaged at the indicated times (hpf); scale bar: 20 μm. (C) Number and size (median and interquartile range are shown) of Golgi objects in the embryo shown in (B) were measured; **, *p* <0.01; ****, *p* < 0.0001; Mann-Whitney test, compared to 8.5 hpf. (D) *Paracentrotus lividus*, (E) *Ciona robusta* and (F) *Branchiostoma lanceolatum* embryos were processed for electron microscopy at the indicated developmental stages; Golgi elements are outlined (isolated stacks in light ochre; linked stacks in light magenta); scale bars: 1 μm. See also Figure S4 and Movie S2.

## Discussion

While the functions played by the ribbon remain unaddressed, indirect evidence suggest that this Golgi configuration must play important functions in cellular physiology. Firstly, the mammalian ribbon disassembles and reassembles during mitosis in proliferating cells; this process involves a finely timed regulation^16^, which is unlikely to have evolved to support the inheritance of a cellular structure with no biological role. Secondly, Golgi “fragmentation”, meaning ribbon disruption, is a morphological phenotype of several human pathologies, among which most notable are neurodegenerative diseases. For instance, in animal models of Amyotrophic Lateral Sclerosis (ALS), Golgi fragmentation precedes phenotypic manifestations^17^. In cellular models of Alzheimer’s, Golgi fragmentation promotes Aβ peptide production^19^, and, in glutamatergic neurons differentiated from induced pluripotent stem cells that carry familial Alzheimer’s disease mutations, Golgi fragmentation is one of the earliest morphological phenotypes detected^23^. From this body of observations, it follows that deciphering which biological functions the Golgi ribbon carries out would not only advance our knowledge of cellular physiology but also help us to better understand the pathological implications of the disruption of this structure. By adopting a comparative perspective that places Golgi architecture in an evolutionary context, here we outline the origins of the ribbon-like organization, propose which molecular effectors may have been responsible for its emergence, and identify its developmentally regulated formation that may signal its function in embryogenesis.

Previously to the present study, the ribbon organization of the Golgi apparatus was thought to be unique to vertebrates. The lack of a centralized Golgi in the cells of *D. melanogaster* and *C. elegans*, two invertebrates widely used in cell biology, may have contributed to cement this view. Nonetheless, works dating to the early 1900’s already showed the presence of ribbon-like Golgi complexes in non-vertebrates^34,35^, and further evidence from electron microscopy analyses of various animal cells accumulated later^36,37,43,83^. Building on this body of literature we sampled species representative of diverse metazoan taxa and show that ribbon-like centralization of Golgi stacks is likely to be a newly evolved character of the ancestor of cnidarians and bilaterians. The frequency with which ribbon-like Golgi complexes are found, both in our morphological analyses and in the literature, supports the generalizations we made on its evolutionary emergence and secondary loss at the level of phyla and superphyla (Figure 1K). Evolutionary conservation of the ribbon-like Golgi configuration in several animal phyla strongly indicates that it must play important functions in their cellular processes. It also begs the question of how did the cells of xenacoelomorphs and ecdysozoans adapt to its secondary loss. Comparative analysis of cellular processes between animals which conserved the ribbon-like Golgi and those which lost it may provide clues regarding the functions this structure is involved in.

Based on experimental evidence from mammalian cells, we propose a plausible molecular mechanism for the appearance of the ribbon-like Golgi organization. GRASP “resurrection” experiments indicate that the self-interacting capability of this protein is ancestral^84^. Bootstrapping on this function, and in the context of cells with multiple Golgi stacks, evolution of GRASP binding by Golgin-45 may have driven ribbon-like emergence in the ancestor of cnidarians/bilaterians. This hypothesis invokes a central role for Golgin-45/GRASP interaction in the evolution and conservation of the mechanism of formation of Golgi ribbons. We focused on Golgin-45 as in mammals it interacts with GRASP55, which is the paralog more similar to the only GRASP present in cnidarians and non-vertebrate bilaterians (Figure S2C). Sequence variations in protein homologs represent natural experiments in mutagenesis. Taking advantage of such experiments done by evolution, we used AlphaFold2 to predict Golgin-45/GRASP binding across holozoans. The results provide support to our hypothesis as they concur with our morphological observations regarding the presence of a ribbon-like Golgi in extant metazoans (Figures 1K and 2E). Whether conspecific Golgin-45 and GRASP proteins do or do not interact as predicted by AlphaFold2 remains to be experimentally tested. It also needs to be established through experiment whether the linked Golgi stacks documented in cnidarians/bilaterians reflect the presence of *bona fide* ribbons; in other words, whether the Golgin-45-dependent spatial orientation of GRASP on Golgi membranes is conducive not only to stack tethering but also to membrane continuity between cisternae of juxtaposed stacks, as observed in mammalian cells^26,27,85^. Mammalian GRASPs are necessary for ribbon formation and GRASP55 interacts with tens of proteins^56,86^. Therefore, it is possible that, by recruiting a network of interactors, oligomerization of correctly oriented GRASP could provide a multivalent molecular platform that directly mediates Golgi stack tethering, and indirectly coordinates the activity of several factors in the formation and maintenance of the Golgi ribbon. Our Golgin-45/GRASP binding hypothesis can be tested in its aforementioned declinations through a comparative approach that takes advantage of the recent addition to lab experimentation of several non-vertebrate animal, such as the cnidarians *Clytia hemisphaerica* and *Nematostella vectensis* or the annelid *Platynereis dumerilii* ^87–89^, and of other established vertebrates and invertebrate experimental organisms.

In eukaryotes, complex multicellularity evolved several times^90^, but non-animal multicellular organisms, such as plants and fungi, display multiple separated Golgi stacks^91–93^. Golgi centralization may thus indicate an evolutionary trajectory with functional requirements specific to cnidarians/bilaterians and divergent from those of basal animals and other multicellular organisms. In this evolutionary context, the question thus arises as to which functions did the ribbon-like Golgi organization evolve to carry out. Were the ribbon, as believed until now, a Golgi configuration restricted to vertebrates, then its functions could have been struck off as specific to this animal lineage. Our findings imply otherwise; at least some of the biological processes the ribbon-like Golgi attends to must be common to all cnidarians/bilaterians. The observation that the ribbon is formed early in embryogenesis (this report and reference^82^) may indicate that, in the context of the whole organism, its first function is linked to development, which would explain why some differentiated mammalian tissues can forgo Golgi ribbons^31–33^. From an evolutionary point of view, it is intriguing to speculate that this may also have been the primordial function of the ribbon-like Golgi. In this hypothetical scenario, the developmental processes of xenacoelomorphs, arthropods and nematodes must have adapted to dispense with the ribbon-like Golgi altogether.

In conclusion, the main finding of the present study is that ribbon centralization into a ribbon-like configuration is common to several animal taxa with well-developed cell types. By the principle of parsimony, we assume that such a situation is explained by a single evolutionary event occurring in the common ancestor of these animals. We expect our report to spark renewed interest in the ribbon configuration of the Golgi apparatus with new studies aimed at testing our hypotheses on its molecular origins and its function in the context of development.

## Supporting information

Sequences of holozoan GRASP protetins

Sequences of holozoan Golgin-45 protetins

## Acknowledgements

We thank Laura Núñez Pons, Filomena Ristoratore, and Periklis Paganos (Stazione Zoologica Anton Dohrn, SZN, Naples, Italy) for providing electron micrographs of sea sponges, of the sea urchin *Strongylocentrotus purpuratus* and helping with the anatomy of the *Ciona intestinalis* larva; Evelyn Houliston (Laboratoire de Biologie du Développement de Villefranche-sur-mer, France) and Mark Terasaki (UConn Health, Farmington, CT, USA) for providing the *Clytia hemisphaerica* electron micrograph; Elisabeth Knust and Michaela Yuan (Max-Planck-Institute of Molecular Cell Biology and Genetics, Dresden, Germany) and Xavier Bailly (Centre National de la Recherche Scientifique & Université Pierre et Marie Curie, CNRS-PMC, Roscoff, France) for their help with *S. roscoffensis* samples; Pedro Martinez Serra (Departament de Genetica, Microbiologia i Estadística, Universitat de Barcelona, Spain) for his valuable suggestions; Martin Lowe (School of Biological Sciences, Division of Molecular & Cellular Function, University of Manchester, UK) and Evelyn Houliston (Laboratoire de Biologie du Développement de Villefranche-sur-mer, France) for their comments on the manuscript. Francesco Ferraro is supported by SZN intramural funding. Emanuele Astoricchio is supported by a PhD fellowship funded by the Stazione Zoologica Anton Dohrn (Open University – Stazione Zoologica Anton Dohrn PhD Program).

## Author contributions

Conceptualization: F.F. Methodology: F.F., M.I.A., S.D., S.L. Investigation: F.F., G.B., S.L., E.A., S.Bo. Resources: S.J., M.K., K.M., V.H., V.B., H.E., S.Be., C.L., J.U-L., B.S., P.B., I.R-T., G.J., E.D. Writing original draft: F.F. Review & Editing: F.F., G.B., S.L., H.E., S.Be, S.D, J.U-L., C.L., I.R-T., B.S., G.J., P.B., M.I.A.. Visualization: F.F. Supervision: F.F.

## Declaration of interests

The authors declare no competing interests.

## Inclusion and diversity statement

The authors support inclusive, diverse, and equitable conduct of research.

## STAR METHODS

### Key Resources Table

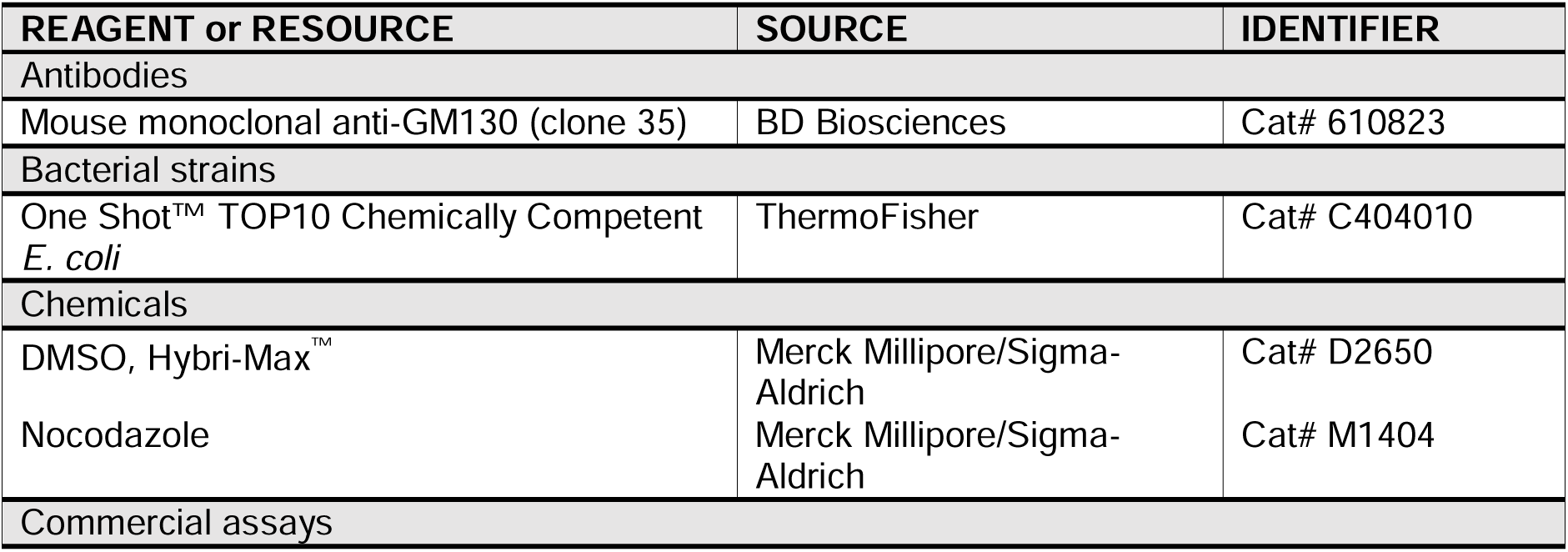
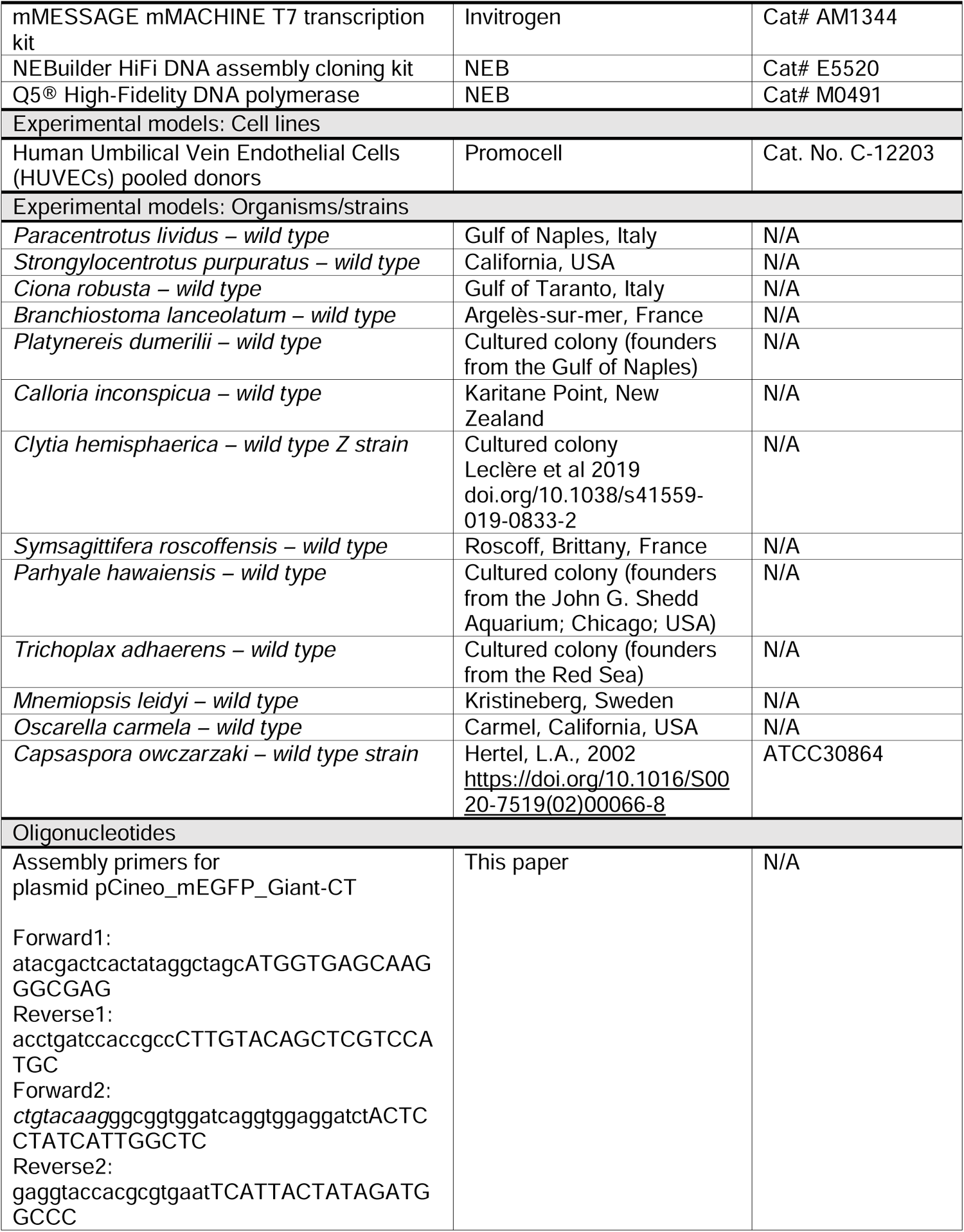
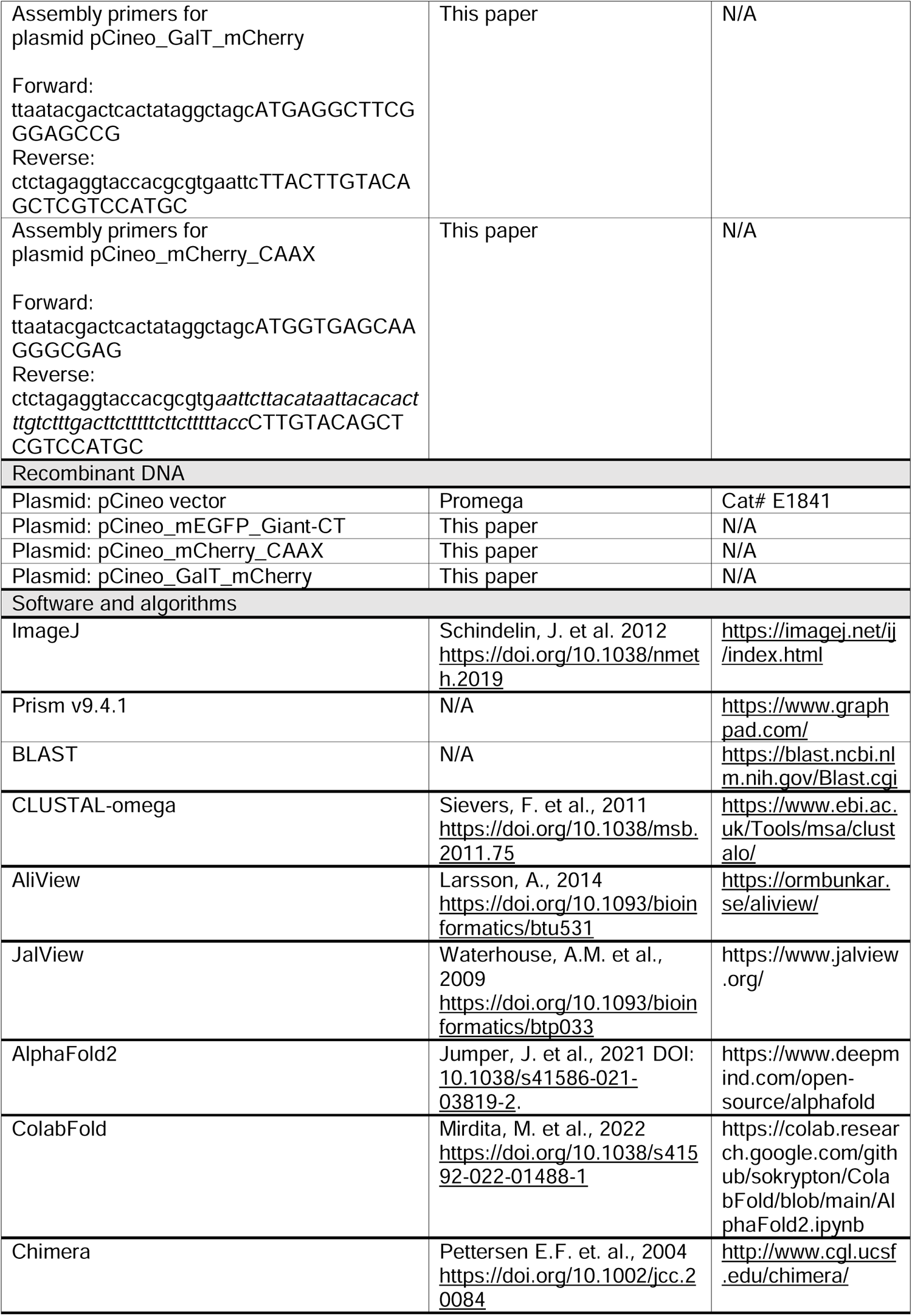
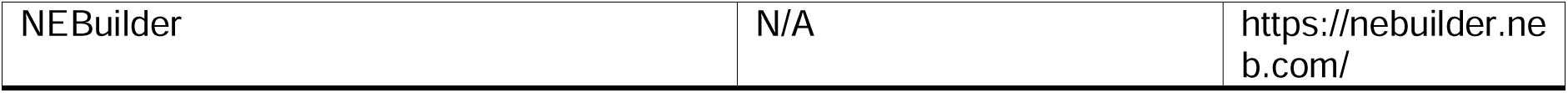

### Resource availability

#### Lead Contact

Further information and request for resources and reagents should be directed to and will be fulfilled by the lead contact, Francesco Ferraro

#### Materials availability

All reagents generated in this study are available from the lead contact upon request.

#### Data and code availability

All data reported in this paper will be shared by the lead contact upon request. This article does not report original code.

### Method details

#### Experimental organisms

Animals were either sourced from the wild or lab cultured. Animal maintenance and treatments to obtain gametes for *in vitro* fertilization have been previously described^44,96–101^. *Parhyale hawaiensis* embryos were a gift by Michalis Averof (Institut de Génomique Fonctionnelle de Lyon, IGFL). *Symsagittifera roscoffensis* juveniles, cultured at 15°C, were processed within 3 days of hatching. The *Capsaspora owczarzaki* ATCC30864 strain, established in 2002^102^, was maintained in modified PYNFH medium (ATCC medium 1034) (https://www.atcc.org/products/327-x).

Details on the sourcing of the organisms used in this study are provided in the Key Resources Table.

#### Cells

Human Umbilical Vein Endothelial Cells (HUVECs), expanded from pools of both sexes acquired from PromoCell, were maintained as described^103^ and used within the 4^th^ passage.

#### Plasmids

Primers were designed using the NEBuilder tool (http://nebuilder.neb.com/). PCR reactions for amplicon generation were carried out with Q5 High-Fidelity DNA Polymerase (NEB). For primer sequences refer to the Key Resources Table. pCineo_mEGFP_Giant-CT (labelled in the figures as mEGFP_Golgi). The plasmid encodes mEGFP in frame with a linker sequence (GGGSGGGS) and the 69 C-terminal amino acids of human Giantin for Golgi membrane targeting. The mEGFP coding sequence was amplified from pmEGFP-N1 vector (Clontech) with primers forward 1 (lower case: pCineo sequence; upper case mEGFP coding sequence) and reverse 1 (lower case: GGGS coding sequence; upper case: mEGFP coding sequence).

The sequence encoding the 69 C-terminal amino acids of human Giantin was amplified from human umbilical vein endothelial cell (HUVEC) cDNA with primers forward 2 (italics: mEGFP coding sequence; lower case: GGGSGGGS linker coding sequence; upper case: Giantin coding sequence) and reverse 2 (lower case: pCineo sequence; upper case: Giantin coding sequence and two stop codons). pCineo_GalT_mCherry. A plasmid (the generous gift of Irina Kaverina, Vanderbilt School of Medicine) encoding the N-terminal 87 amino acids of galactosyl-transferase (GalT), which confer Golgi localization, in frame with mCherry was used as template to amplify the GalT_mCherry coding sequence using primers forward (lower case: pCineo sequence; upper case: GatT coding sequence) and reverse (lower case: pCineo sequence; upper case: GatT coding sequence). pCineo_mCherry_CAAX (labelled in the figures as mCherry_PM). The sequence encoding mCherry in frame with the polybasic sequence and CAAX motif of human K-Ras (GKKKKKKSKTKCVIM) for targeting to the plasma membrane was generated by amplification of mCherry using the pmCherry-N1 (Clontech) plasmid as template and the following primers: forward (lower case: pCineo sequence; upper case: mCherry coding sequence) and reverse (lower case: pCineo sequence; italics: polybasic plus CAAX motif and stop codon coding sequence; upper case: mCherry coding sequence).

Amplicons and pCineo plasmid (linearized by NheI/EcoRI digestion) were assembled using the NEBuilder HiFi DNA assembly cloning kit (NEB) following the manufacturer instructions. Correct sequences were verified by Sanger sequencing.

#### *In vitro* mRNA transcription and microinjections

Plasmids were linearized by digestion with NotI, a unique restriction site in the pCineo vector located downstream of the cloned sequences. One microgram of each linearized plasmid was used as template for *in vitro* transcription. Purified mRNAs were resuspended in DEPC-MilliQ water, their concentration measured, and their quality checked by agarose gel electrophoresis. mRNAs were aliquoted and stored at – 80°C until used.

Sea urchin eggs’ jelly coat was dissolved by a short wash in acidic filtered sea water (1.5 mM citric acid in 0.22 μm filtered sea water, FSW). De-jellied eggs were then immobilized on 60 mm plastic dish lids pre-treated with 1% protamine sulphate (Merck, Sigma-Aldrich, P4380) in FSW. Eggs were then washed with FSW containing sodium para-amino benzoate (Sigma-Aldrich, A6928; 0.05% in FSW) to prevent hardening of the fertilization envelope. *In vitro* transcribed mRNAs were diluted to a final concentration of 300-500 ng/μL in 120 mM KCl/DEPC-water. Four to five pL of diluted mRNAs were injected per embryo, immediately after fertilization. Embryos were allowed to develop at 18°C.

#### Confocal microscopy

*Paracentrotus lividus*. At the indicated times post-fertilization, embryo development was stopped by incubation with 0.2% paraformaldehyde in FSW, which kills the embryos while preserving mEGFP and mCherry fluorescence. Imaging was carried out within 16 h of formaldehyde treatment. Embryos laid on glass-bottom dishes containing FSW were imaged with an inverted 25x (NA 0.8) water immersion objective, using a Zeiss LSM700 system. Image stacks (z-step 1 μm) were acquired. Only one third to one half of the embryo volumes could be imaged at early stages, due to the opacity of yolk granules. At later stages (prism and pluteus) embryos were transparent and their whole volume was imaged.

For live imaging experiments, eggs were laid in FWS containing glass-bottom dishes pre-treated with protamine, fertilized, and then immediately microinjected with fluorescent reporter encoding mRNAs. Imaging was carried out as described above. Image stacks (z-step 1 μm) were acquired at 15 min intervals. Higher magnification imaging of embryos was carried out on mEGFP_Giant-CT (mEGFP_Golgi) microinjected embryos using a 40x (NA 1.10) water immersion objective with a Leica SP8 confocal system. For presentation purposes, contrast-enhancement and gaussian-blur filtering were carried out (ImageJ) to the images shown. *HUVECs*. Cells were seeded on gelatin-coated 96-well plates (Nunclon surface©, NUNC) at 15.000 cells/well and grown in HGM medium for 24 h. After rinsing with fresh medium, cells were fed HGM containing 0.1% (vol:vol) DMSO (control treatment) or 33 µM (10 mg/mL) Nocodazole and incubated for hours before fixation with 4% formaldehyde in phosphate-buffered saline (PBS) for 10 minutes at RT. Fixed cells were permeabilized with 0.2% TX-100 (Merck, Sigma-Aldrich) in PBS for 10 min (RT) and then blocked with 5% BSA (Merck, Sigma-Aldrich) in PBS for 30 min (RT). The Golgi apparatus was immuno-labeled with an antibody raised against the Golgi marker GM130 (BD Biosciences), followed by incubation with Alexa Fluor 488 conjugated anti-mouse antibody (Life Technologies); primary and secondary antibodies were diluted in 1% BSA/0.02% TX-100/PBS. Nuclei were counterstained with Hoechst 33342 (Life Technologies), diluted in PBS, and images acquired using an Opera High Content Screening System (Perkin Elmer) through a 40x air objective (NA 0.6). Exclusively for presentation purposes, the confocal images of sea urchin embryos were subjected to gaussian blurring (ImageJ) of the Golgi and plasma membrane channels.

#### Image analysis

Golgi objects from confocal images were analyzed with ImageJ (https://imagej.nih.gov/ij/). The Golgi channel (8-bit) was selected, maximum intensity projection images generated and processed as follows.

Time course (Figure 3A). All images were subjected to background subtraction. Small Golgi objects observed at 2, 4 and 6 hpf were identified with the “find maxima” command and separated from each other by segmentation. The images of all time points were then subjected to thresholding and transformed into binary images. Golgi object number and size were then counted with the “analyze particles” command (area range was set at 0.25 – infinite μm^2^). Three embryos per time point were analyzed. Time-lapse (Figure 3B). Image threshold was set automatically. At early time points, slight adjustments were done to correctly capture the size of most Golgi objects. For later time points, default threshold values were sufficient to correctly outline the size Golgi objects. After transformation into binary images, object number and size were measured as described above. Numerical results were processed with Prism (Graphpad) for graph plotting and statistical analysis.

#### Electron microscopy

*Paracentrotus lividus*, *Branchiostoma lanceolatum* and *Ciona robusta* samples, maintained at 18°C, were collected at the indicated developmental stages and fixed at 4°C in 2% glutaraldehyde in filtered sea water (FSW). After fixation samples were first rinsed in FSW (6 x 10 min), then in Milli-Q water (3 x 10 min) and post-fixed with 1% osmium tetroxide and 1.5% potassium ferrocyanide (1 h, 4°C). Samples were then rinsed five times with Milli-Q water, dehydrated in a graded ethanol series, further substituted by propylene oxide and embedded in Epon 812 (TAAB, TAAB Laboratories Equipment Ltd, Berkshire, UK). Resin blocks were sectioned with a Ultracut UCT ultramicrotome (Leica, Vienna, Austria). Sections were placed on nickel grids and observed with a Zeiss LEO 912AB TEM (Zeiss, Oberkochen, Germany).

##### Calloria inconspicua

Three-lobed larvae were initially fixed in 2.5% glutardialdehyde buffered with 0.1 M sodium cacodylate solution (60 min at 5°C). A tiny amount ruthenium red solution was added to stain the extracellular matrix. Repeated rinsing in 0.1 M sodium cacodylate buffer was followed by post-fixation in 1% osmium tetroxide solution buffered with 0.1 M sodium cacodylate (40 min at 4°C). Dehydration with an acetone series and propylene oxide led to embedding in Araldite. Resin blocks were polymerized at 60°C for 48 hours. Ultrathin serial sections (70 nm) were cut on a Reichert Ultracut E microtome, placed on formvar-coated copper slot grids, and automatically stained with uranyl acetate and lead citrate in a LKB Ultrostainer. The sections were examined in Zeiss EM 10B and Zeiss EM 900 transmission electron microscopes.

##### Parhyale hawaiensis

Embryos were pre-fixed in 2.5% glutardialdehyde, 2% paraformaldehyde, 2% sucrose in sodium cacodylate buffer 0.1 M (SC buffer) overnight at 4°C. After several rinses in SC buffer at room temperature specimens were postfixed in 1% OsO4 in 0.1 M SC Buffer (2 hrs, room temperature), washed in SC buffer (1 hr) and dehydrated in an ethanol series. Ethanol-preserved specimens were sent to Berlin, transferred to 100% acetone and propylene oxide and subsequently embedded in araldite. Ultrathin sections were cut on a Leica EM UC7, stained with Plano uranyl acetate replacement stain (UAR-EMS) and lead citrate and investigated in a LEO EM 906.

*Strongylocentrotus purpuratus, Platynereis dumerilii*, *Mnemiopsis leidyi*, *Oscarella carmela* and *Capsaspora owczarzaki* samples were high-pressure frozen, freeze substituted and processed as described^44,45,47, 104–106^.

##### Trichoplax adhaerens

Animals, alive of pre-fixed, were high-pressure frozen/freeze substituted and embedded in Epon. Sections (70 nm) were cut with using a Leica Ultracut UCT ultramicrotome.

##### Symsagittifera roscoffensis

Animals were processed within three days of hatching. The head of a hatchling was processed by high-pressure freezing. Freeze substitution was carried out in a solution of 1% osmium tetroxide and 0.1% uranyl acetate in acetone. A Leica Ultracut UCT was used to generate 60–80 nm sections, which were poststained in a 2% uranyl acetate/lead citrate solution and transferred to formvar-coated slit grids. Sections were imaged with a Tecnai 12 Biotwin TEM, using a fast-scan F214A CCD camera controlled by the SerialEM software (Boulder Lab). Digital image stacks were imported into the TrakEM2 package.

##### Clytia hemisphaerica

Individual ovaries were high-pressure frozen with a Wohlwend Compact 03 high-pressure freezing machine (http://www.wohlwend-hpf.ch) using sea water as the freezing medium and then transferred to a frozen solution of 2% osmium in acetone under liquid nitrogen. The ovaries were freeze-substituted in a Leica AFS2 freeze-substitution machine (https://www.leica-microsystems.com) using the following program: −90°C for 18 hours, −90°C to −30°C with a slope of 5°C/hour, - 30°C for 12 hours, −30°C to 0°C with a slope of 5°C/hour. Samples were removed from the AFS chamber and allowed to reach room temperature. This was followed by 5 acetone washes for 5 minutes each. Ovaries were infiltrated with Polybed resin in a series of steps as follows: 1:3 resin to acetone overnight, 1:1 resin to acetone for 6 hours, 3:1 resin to acetone overnight, 100% resin for 6 hours followed by embedment in molds in fresh 100% resin and curation at 60°C for 2 days. Polymerized samples were then trimmed using an ultramicrotome to get the entire cross-section of the ovary. Serial 60 nm sections were collected using an Automated Tape-Collecting Ultramicrotome, mapped, and imaged with a Zeiss Sigma FE-SEM as described previously^107^.

#### Homology search

Canonical human GRASP (GRASP65 and GRASP55) and Golgin-45 amino acid sequences (Data S1 and S2) were used as initial queries. Homologs were searched in the target species using Basic Local Alignment Search Tool, BLAST, (BLASTp and TBLASTn) in available databases (Uniprot, NCBI, Ensembl). For specific target species, the search was carried out in dedicated databases (*Amphiura filiformis*: http://www.echinonet.org.uk/blast/; *Mnemiopsis leidyi*: https://research.nhgri.nih.gov/mnemiopsis/sequenceserver/; *Nematostella vectensis*: http://marimba.obs-vlfr.fr/blast; unicellular holozoans: https://protists.ensembl.org). Target genomes and, whenerver available, transcritpomes were interrogated. Hits with the lowest E-value and highest query coverage were selected as candidate homologs and validated when by reverse BLAST on the human proteome the query was retrieved as the highest scoring. If this approach did not return a hit, homologs of evolutionarily closer species were used as queries. Further validation of homology was obtained by subjecting the hits to sequence and structural analysis with InterProScan (https://www.ebi.ac.uk/interpro/search/sequence/) and by multiple sequence alignment with AliView and JalView to verify regions of sequence similarity.

#### Structure modelling

Models of complexes between conspecific GRASP/Golgin-45 pairs were built with the Colab implementation of AlphaFold2^76^, using MMseqs2 to generate multiple sequences alignments^108^. To obtain reliable predictions of the protein-peptide complexes, AlphaFold-Multimer version v2 was used, with 12 recycles for the generation of each model^109^. Complexes were built without the use of structural templates and without Amber refinement as this step does not introduce substantial improvement, while significantly increasing computational time.

#### Note about the phylogenetic trees

Whether sponges or ctenophores or placozoans are the sister group to all other animals remains an unsettled issue^110–120^; for this reason we drew the holozoan tree of life as a polytomy of these three taxa in Figures 1K, 2E and in the graphical abstract. The animal silhouettes used in the graphical abstract were obtained from the public domain (http://phylopic.org), when not covered by copyright, or drawn by F.F.

## Notes

### Competing Interest Statement

The authors have declared no competing interest.

### Summary of Updates

This version includes an Introduction and the order of the presented data was changed with respect the previous version.

