## Supplemental Information for "Evolution of the ribbon-like organization of the Golgi apparatus in animal cells"

**List of Supplemental information**

- **Supplemental Figures**

Figures S1-S4

- **Supplemental Data**

Data S1-S2

- **Supplemental Movies**

Movies S1-S2

- **Supplemental Results and Discussion**
- **Supplemental References**

**Legends**

**Figure S1. Additional examples of Golgi structure in holozoans**. Related to Figure 1. (A) Golgi stack array in secretory cells of *Symsagittifera roscoffensis*. (B) A three-day-old *Platynereis* *dumerilii* larva. Serial sections (40 nm each), labelled starting from 1, are shown; in the region of interest, separated stacks (labelled a, b, c and d) in section 1 are seen to merge (c + d and then b + c + d) into a ribbon while progressing through the sections. (C) Comb cells of *Mnemiopsis leidyi*. (D-E) Two cell types of *Trichoplax adhaerens*. Scale bars: 1 μm. (F) Table summarizing Golgi organization in species and cell types discussed in this report.

**Figure S2. Structural features of holozoan GRASP and Golgin-45 proteins**. Related to Figure 2. (A) Cartoon of domain structure of mammalian GRASPs. The evolutionarily conserved GRASP domain is formed by a tandem of atypical PDZ domains, followed by a C-terminal region, which in mammals is serine/proline-rich (SPR) and whose post-translational modifications modulate GRASP activity. (B) Size, in amino acids, of the GRASP domains and C-terminal regions of the holozoan GRASP sequences (Data S1) were plotted; bars indicate median size. (C) Pairwise amino acid identity of holozoan GRASP domains plotted as a heat map. Vertebrate duplication into GRASP55 and GRASP65 paralogs occurred with the evolution of jawed vertebrates. In vertebrates, GRASP55 paralogs (green outlines) are more similar to bilaterian single GRASPs than GRASP65 paralogs (red outlines); percent identity values for pairwise comparisons were obtained with CLUSTAL omega. (D) Structure of the GRASP domain (gold) of mouse GRASP55 in complex with the C-terminal residues of mouse Golgin-45 (ball and stick); PDB accession number 5H3J. PDZ1 and PDZ2 are indicated by color coded circles. Golgin-45 residues important for the interaction are highlighted; green, the stretch of residues interacting with the groove formed by PDZ1 and PDZ2 domains (arrows indicate F390 and N391, which are discussed in the “Structure modelling” section of the Method details and Figure S3); magenta, the two cysteines involved in Zinc-like finger formation; light blue, the PDZ-binding motiI(E) Multiple sequence alignment of the C-termini of the holozoan Golgin-45 homologs. The residues corresponding to the binding features are color coded as in Figure S2D. Insertions (maroon highlight) and deletions (dashed boxes) are indicated relative to the mouse sequence; variations mostly affect the groove-binding sequence.

**Figure S3. Additional AlphaFold2 models**. Related to Figure 2. (A) Model of Golgin-45/GRASP complex of the only xenacoelomorph species with a Golgin-45 gene (Data S2). (B) Models of mouse Golgin-45 mutants in complex with mouse GRASP domain; color-coded arrows indicate altered binding features as detailed in Figure 3D.

**Figure S4. Golgi dynamics in the sea urchin embryo**. Related to Figure 3. (A) The fluorescent reporter used in this study, mEGFP_Golgi, co-localizes with the widely used Golgi reporter GalT_mCherry in the sea urchin (*P. lividus*) early gastrula; scale bar: 20 μm. (B) Quantification of Golgi object size (n = 3 embryos) from the time-course experiment shown in Figure 1A; ****, p < 0.0001 (Mann-Whitney test). (C) *Paracentrotus lividus* embryos expressing the mEGFP_Golgi reporter imaged at the indicated stages; scale bar: 50 μm. (D) Golgi apparatus imaging of a 15 hpf *Paracentrotus lividus* embryo. A single focal plane acquired with a 40x water immersion objective is shown; scale bar: 5 μm. I Golgi stacks a, b and c are seen establishing connections across serial sections (numbered in black), of a blastocoel cell of the sea urchin *Strongylocentrotus purpuratus* pluteus; scale bar: 1 μm. (F) Golgi disassembly/reassembly during mitosis in the *Paracentrotus lividus* embryo; image series (left to right, 15 min acquisition interval); scale bar: 5 μm. (G-H) Treatment with the microtubule depolymerizing compound nocodazole induces ribbon unlinking into constituent Golgi stacks in human umbilical vein endothelial cells, HUVECs, and sea urchin embryos; magnifications of insets are shown; scale bars: 10 μm and 20 μm (HUVECs and sea urchin, respectively).

**Data S1. Holozoan GRASP homologs**. Sequences of selected holozoan GRASP proteins are reported. GRASP domains, defined as residues number 1 to the fifth following the invariant motif His-Arg-Iso-Pro at the end of the second PDZ domain, are highlighted in bold; the conserved glycine residues in position 2 are highlighted in red.

**Data S2. Holozoan Golgin-45 homologs**. Sequences of selected holozoan Golgin-45 proteins are reported. InterProScan analysis identified all the sequences as Golgin-45/BLZF1-like. Among xenacoelomorphs, a Golgin-45 homolog was found only in the species *Hofstenia miamia*.

**Movie S1. Golgi ribbon in a glial cell of the three-day-old *Platynereis dumerilii* larva**. Related to Figure 1. Image series of volume EM. The series includes the image shown in Figure 1D.

**Movie S2. Golgi clustering in the sea urchin embryo**. Related to Figure 3. Time-lapse microscopy of Golgi element dynamics in a *Paracentrotus lividus* embryo (shown as image series in Figure 3B). Maximum intensity projection of image stacks acquired at 15 min intervals between 7h:30m and 10h:15m post-fertilization are shown.

**Supplemental Results and Discussion**

**Interpretation and predictive power of AlphaFold2 models**. The introduction of AlphaFold^1^ and its subsequent evolutions produced a revolution in structural biology, allowing the generation of structure models of unprecedented accuracy. Recent benchmark studies have demonstrated that the predictive power of AF2 extends beyond the production of the mere structural models, yielding accurate results also for protein-protein and protein-peptide complexes, even when they imply conformational changes, and providing reliable hints on the effect of missense mutations^2^. In the case of protein-peptide complexes, models with higher confidence can be obtained by increasing the number of recycles during model generation, provided that a sufficient number of sequences are detected during the generation of the multiple sequence alignments (MSAs)^3,4^. In general, AF2 predicted models are evaluated and ranked based on a per residue score, the predicted local distance difference test (pLDDT). This value provides a measure, from a minimum of 0 to a maximum of 100, of the agreement between the prediction and experimental structures. Models or regions within them with an average pLDDT ≥ 70 are generally considered reliable^5^. At the same time, it has been observed that low pLDDT values are indicative of intrinsically disordered regions and highly flexible stretches within proteins^6,7^. Therefore, stable complexes, in which the binding partners have reduced mobility with respect to each other, are typically modelled with higher pLDDT scores. With all these considerations in mind, we built models for representative pairs of Golgin-45/GRASP from different species; the GRASPs from all species were modeled with extremely high confidence (average pLDDT ≥ 90), whereas variable results were obtained for the C-terminal peptides of the Golgin-45 proteins. In particular, peptides lacking the PDZ-binding motif (e.g., *D. melanogaster*) and/or the cysteines for Zn-finger formation (e.g., *M. leidyi*) could not adopt the binding conformation and were associated to very low pLDDT values. In general, lower pLDDT values characterized the residues interacting with the groove. While these scores could partially arise from low sequence coverage in the MSAs, they may also be indicative of higher mobility of said regions, i.e., absence of interaction with the groove and, in some cases, complete displacement of the Golgin-45 peptide.

**Role of groove residues in Golgin-45/GRASP interaction**. To validate the structural conclusions derived from the crystal structure of the mouse GRASP domain (of GRASP55) in complex with the Golgin-45 C-terminus, the authors of that study performed binding assays of protein mutants by pulldown experiments and isothermal titration calorimetry^8^. Mutation of the last Golgin-45 residue (I403R), which disrupts the PDZ-binding motif abolished binding to the GRASP domain; this was also the case when the cysteines involved in the Zinc finger formation were mutated (C393A, C3956A)^8^. These results were correctly reproduced in AF2 models. In fact, as all modeled complexes have identical levels of sequence coverage, pLDDT decreases compared to the reference structure can in this case be ascribed to increased flexibility, decreased interaction, and reduced binding, and the displacement of the Golgin-45 peptide from its binding site is clearly visible in the structures obtained (Figure S3B). With respect to the interaction with the GRASP groove, the authors of the above study carried out binding assays with two Golgin-45 mutants, F390A and N391A, which did not abolish its interaction with the GRASP domain^8^. From these results they concluded that the groove-binding residues of Golgin-45 play little or no role in GRASP interaction. However, the mutation of F390, which in the crystal structure is buried in the hydrophobic groove of the GRASP domain, to alanine is too conservative and should not impact groove binding. At the same time, the side chain of N391 in the crystal structure faces outwards from the GRASP groove, therefore also the mutation N391A is expected to have low impact on the interaction (Figure S2D). We modelled F390A and N391A mutations with AlphaFold2, finding, as expected, that they do not significantly alter the conformation of the Golgin-45 C-terminus and of the complex (Figure S3B). We conclude that these mutations are not ideal to assess the role of the interaction with the groove in the overall binding. For instance, replacement of F390 with a charged residue (F390R or F390E), which cannot be accommodated in the hydrophobic GRASP pocket, produced models devoid of the peptide/groove interaction and might be more indicated to experimentally validate the role of this interaction (Figure S3B). Of note, substitution of a charged residue in place of hydrophobic I388 (I388R) while making the region more flexible is still predicted to be able to fit the groove (Figure S3B), suggesting that some of the groove-inserting residues may be less important than others (e.g., F390) for the interaction with the GRASP domain. Although our modeling results may indicate that, at least in deuterostomes (with their highly conserved Golgin-45 and GRASP sequences), the Golgin-45 residues projecting into the GRASP groove increase binding stability, their actual contribution awaits experimental confirmation. Therefore, we considered stable binding between holozoan Golgin-45 and GRASP to occur when in their models both PDZ-binding motif interaction and Zinc finger formation were detected. It is worth noting that the conclusions regarding the appearance of Golgin-45/GRASP during evolution (Figure 2E) would not substantially change even in the case the Golgin-45 groove-interacting residues were shown to be required for GRASP binding (Figures 2D and S3A, green arrow).

**Supplemental Refences**

1.	Jumper, J., Evans, R., Pritzel, A., Green, T., Figurnov, M., Ronneberger, O., Tunyasuvunakool, K., Bates, R., Zidek, A., Potapenko, A., et al. (2021). Highly accurate protein structure prediction with AlphaFold. Nature *596*, 583-589. 10.1038/s41586-021-03819-2.

2.	Akdel, M., Pires, D.E.V., Pardo, E.P., Janes, J., Zalevsky, A.O., Meszaros, B., Bryant, P., Good, L.L., Laskowski, R.A., Pozzati, G., et al. (2022). A structural biology community assessment of AlphaFold2 applications. Nat Struct Mol Biol *29*, 1056-1067. 10.1038/s41594-022-00849-w.

3.	Tsaban, T., Varga, J.K., Avraham, O., Ben-Aharon, Z., Khramushin, A., and Schueler-Furman, O. (2022). Harnessing protein folding neural networks for peptide-protein docking. Nature communications *13*, 176. 10.1038/s41467-021-27838-9.

4.	Johansson-Akhe, I., and Wallner, B. (2022). Improving peptide-protein docking with AlphaFold-Multimer using forced sampling. Front Bioinform *2*, 959160. 10.3389/fbinf.2022.959160.

5.	Tunyasuvunakool, K., Adler, J., Wu, Z., Green, T., Zielinski, M., Zidek, A., Bridgland, A., Cowie, A., Meyer, C., Laydon, A., et al. (2021). Highly accurate protein structure prediction for the human proteome. Nature *596*, 590-596. 10.1038/s41586-021-03828-1.

6.	Guo, H.B., Perminov, A., Bekele, S., Kedziora, G., Farajollahi, S., Varaljay, V., Hinkle, K., Molinero, V., Meister, K., Hung, C., et al. (2022). AlphaFold2 models indicate that protein sequence determines both structure and dynamics. Sci Rep *12*, 10696. 10.1038/s41598-022-14382-9.

7.	Ruff, K.M., and Pappu, R.V. (2021). AlphaFold and Implications for Intrinsically Disordered Proteins. Journal of molecular biology *433*, 167208. 10.1016/j.jmb.2021.167208.

8.	Zhao, J., Li, B., Huang, X., Morelli, X., and Shi, N. (2017). Structural Basis for the Interaction between Golgi Reassembly-stacking Protein GRASP55 and Golgin45. The Journal of biological chemistry *292*, 2956-2965. 10.1074/jbc.M116.765990.
