## Supplemental Figures S1-S4 for "Evolution of the ribbon-like organization of the Golgi apparatus in animal cells"

### Figure S1

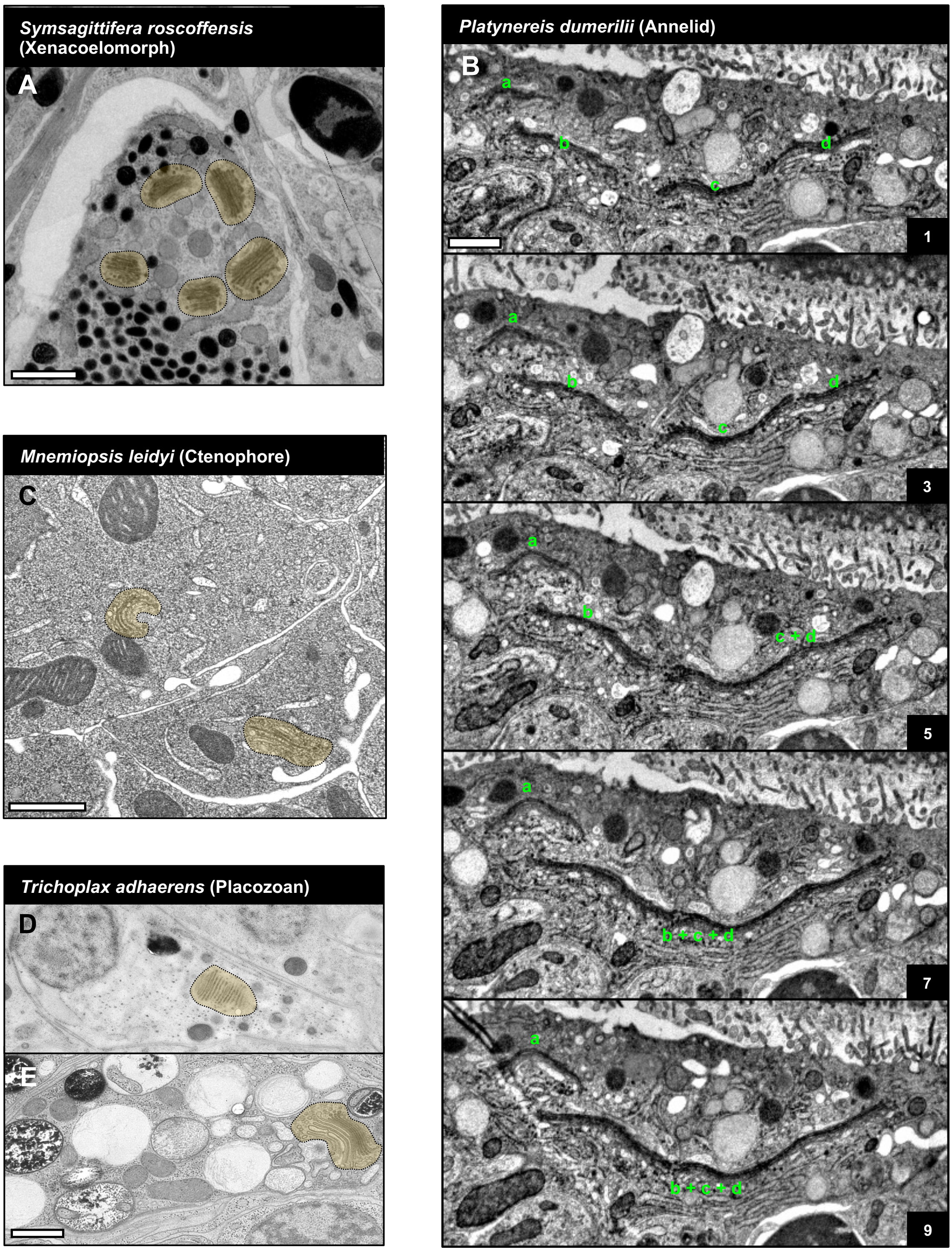


**F**

| Species | Taxon | Multiple Golgi stacks per cell | Presence of ribbon-like Golgi | Source of morphological data | Tissues/cell types |
|---|---|---|---|---|---|
| *C. robusta* | Tunicates | + | + | This paper | epidermal cells of larva |
| *B. lanceolatum* | Cephalocordates | + | + | This paper | ectodermal cells of gastrula |
| *S. purpuratus* | Echinoderms | + | + | This paper | blastocoelar cells of pluteus larva |
| *P. lividus* | Echinoderms | + | + | This paper | all cells from pre-hatching blastula to pluteus larva |
| *L. variegatus* | Echinoderms | + | + | Literature | all cells in pre-hatching blastula |
| *L. pictus* | Echinoderms | + | + | Literature | all cells in blastula and prism stage |
| *D. melanogaster* | Arthropods | + | – | Literature | all cell types (larva and adult) and cell lines |
| *A. mellifera* | Arthropods | + | – | Literature | Trophocyte |
| *A. pisum* | Arthropods | + | – | Literature | Mycetocytes |
| *A. albopictus* | Arthropods | + | – | Literature | cell line |
| *P. hawaiensis* | Arthropods | + | – | This paper | all cell types (adult) |
| *C. elegans* | nematodes | + | – | Literature | all cell types |
| *C. inconspicua* | Brachiopods | + | – | This paper | epidermal cells of 3-lobe larva |
| *P. vivipara* | Mollusks | + | + | Literature | spermatocytes |
| *H. pomatia* | Mollusks | + | + | Literature | multified gland cells |
| *H. aspersa* | Mollusks | + | + | Literature | early spermatocytes |
| *P. dumerilii* | Annelids | + | + | This paper | several cell types of 3-day-old larva |
| *Lumbricus (unreported species)* | Annelids | + | + | Literature | epithelial cells and neurons of adult |
| *S. roscoffensis* | Xenacoelomorphs | + | + | This paper | secretory granule producing cells |
| *C. hemisphaerica* | Cnidarians | + | + | This paper | gastrodermal cells of adult |
| *M. leidyi* | Ctenophores | + | – | This paper | epithelial and comb cells |
| *T. adhaerens* | Placozoans | – | N/A | This paper | all cell types |
| *H. hongkongensis* | Placozoans | – | N/A | Literature | all cell types |
| *O. carmela* | Sponges | – | N/A | Literature and this paper | all cell types |
| *E. fluviatilis* | Sponges | + | – | Literature | spongocyte |
| *S. rosetta* | Choanoflagellates | – | N/A | Literature | N/A |
| *C. owczarzaki* | Filasterians | – | N/A | This paper | N/A |

### Figure S2

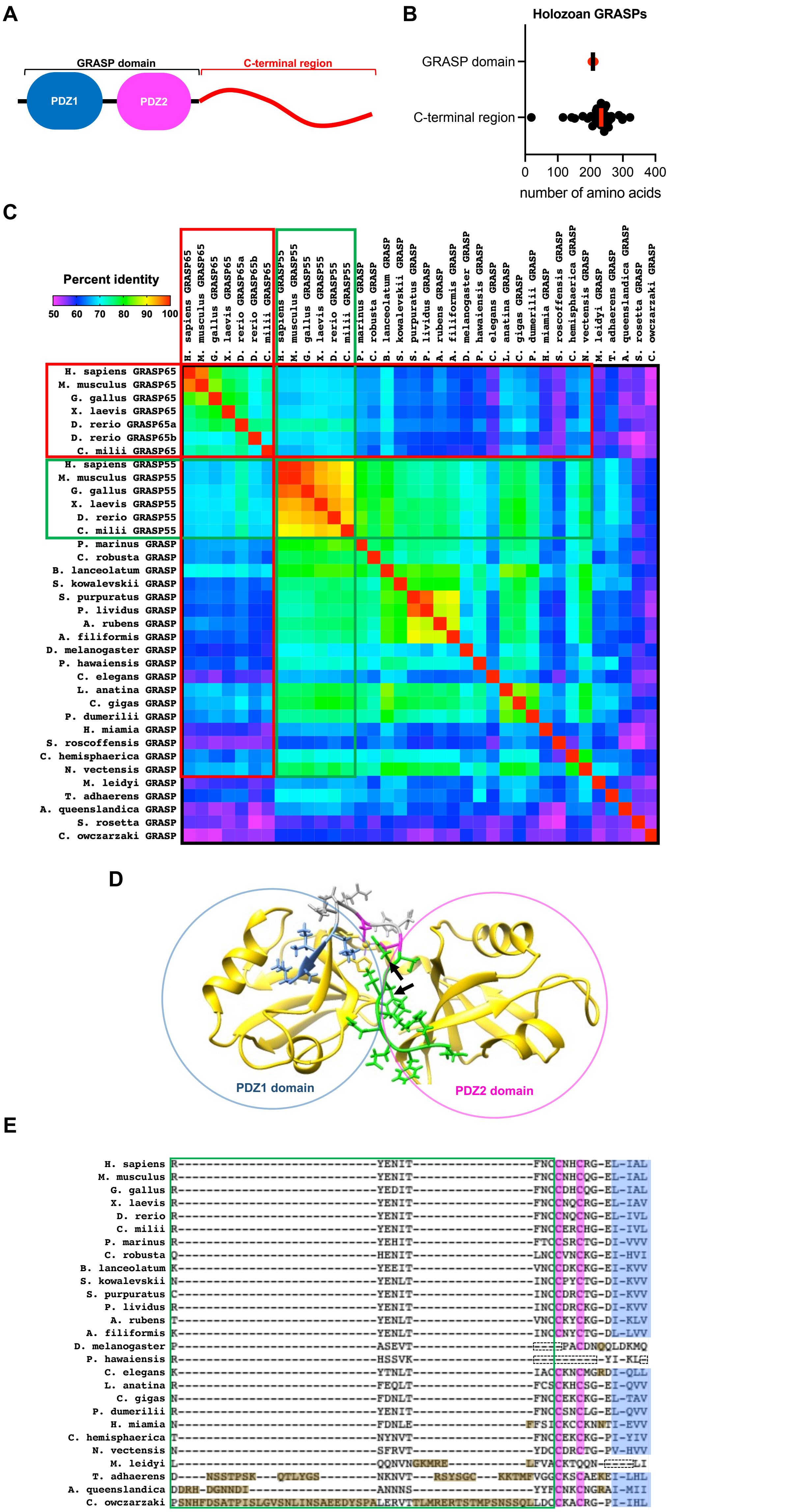
A
GRASP domain
C-terminal region
PDZ1
PDZ2
B
Holozoan GRASPs
GRASP domain
C-terminal region
0
100
200
300
400
number of amino acids
C
Percent identity
50
60
70
80
90
100
H. sapiens GRASP65
M. musculus GRASP65
G. gallus GRASP65
X. laevis GRASP65
D. rerio GRASP65a
D. rerio GRASP65b
C. milii GRASP65
H. sapiens GRASP55
M. musculus GRASP55
G. gallus GRASP55
X. laevis GRASP55
D. rerio GRASP55
C. milii GRASP55
P. marinus GRASP
C. robusta GRASP
B. lanceolatum GRASP
S. kowalevskii GRASP
S. purpuratus GRASP
P. lividus GRASP
A. rubens GRASP
A. filiformis GRASP
D. melanogaster GRASP
P. hawaiensis GRASP
C. elegans GRASP
L. anatina GRASP
C. gigas GRASP
P. dumerilii GRASP
H. miamia GRASP
S. roscoffensis GRASP
C. hemisphaerica GRASP
N. vectensis GRASP
M. leidyi GRASP
T. adhaerens GRASP
A. queenslandica GRASP
S. rosetta GRASP
C. owczarzaki GRASP
D
PDZ1 domain
PDZ2 domain
E
H. sapiens
M. musculus
G. gallus
X. laevis
D. rerio
C. milii
P. marinus
C. robusta
B. lanceolatum
S. kowalevskii
S. purpuratus
P. lividus
A. rubens
A. filiformis
D. melanogaster
P. hawaiensis
C. elegans
L. anatina
C. gigas
P. dumerilii
H. miamia
C. hemisphaerica
N. vectensis
M. leidyi
T. adhaerens
A. queenslandica
C. owczarzaki

### Figure S3

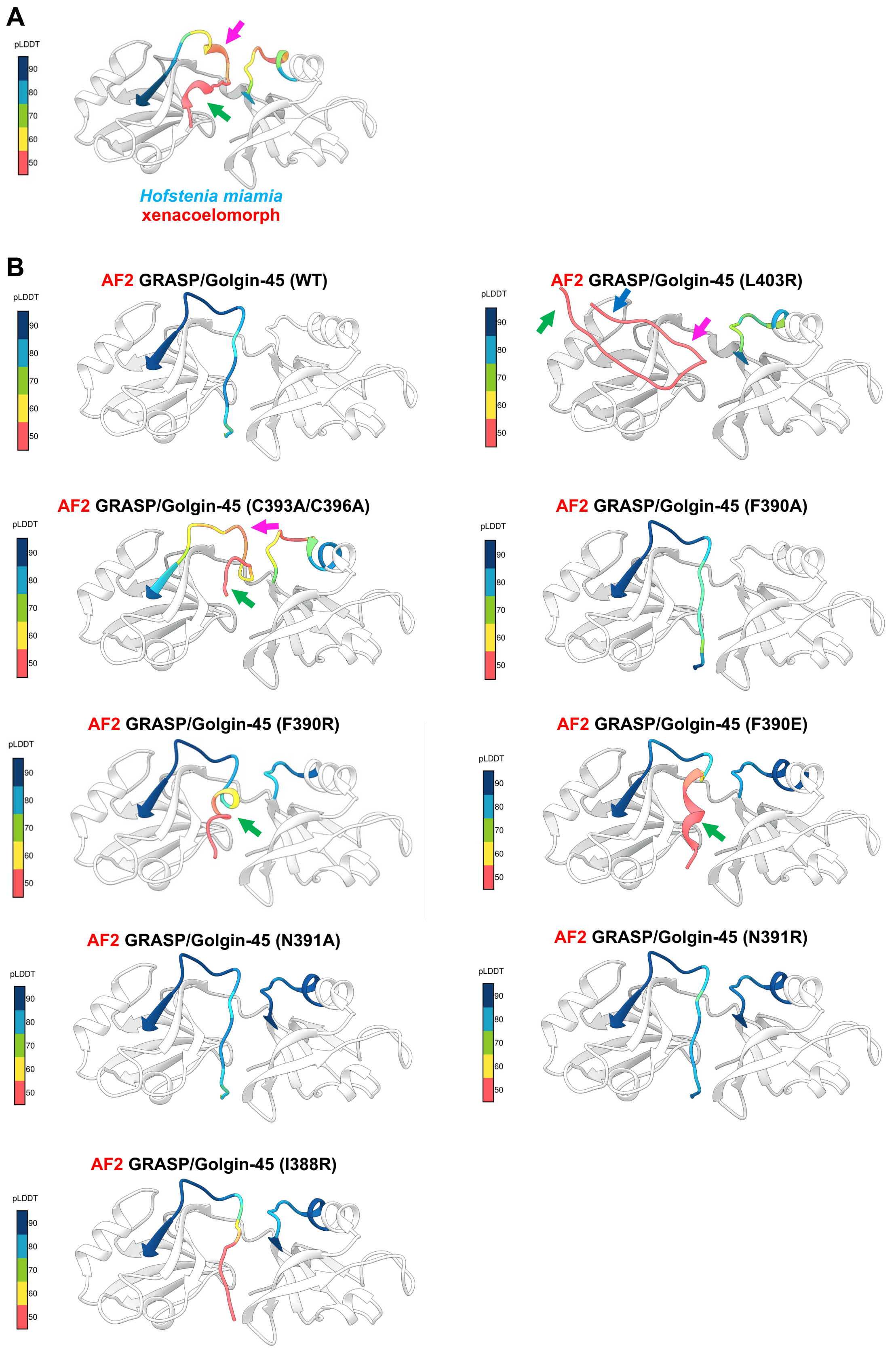
A
pLDDT
90
80
70
60
50
Hofstenia miamia
xenacoelomorph
B
AF2 GRASP/Golgin-45 (WT)
AF2 GRASP/Golgin-45 (L403R)
AF2 GRASP/Golgin-45 (C393A/C396A)
AF2 GRASP/Golgin-45 (F390A)
AF2 GRASP/Golgin-45 (F390R)
AF2 GRASP/Golgin-45 (F390E)
AF2 GRASP/Golgin-45 (N391A)
AF2 GRASP/Golgin-45 (N391R)
AF2 GRASP/Golgin-45 (I388R)

### Figure S4

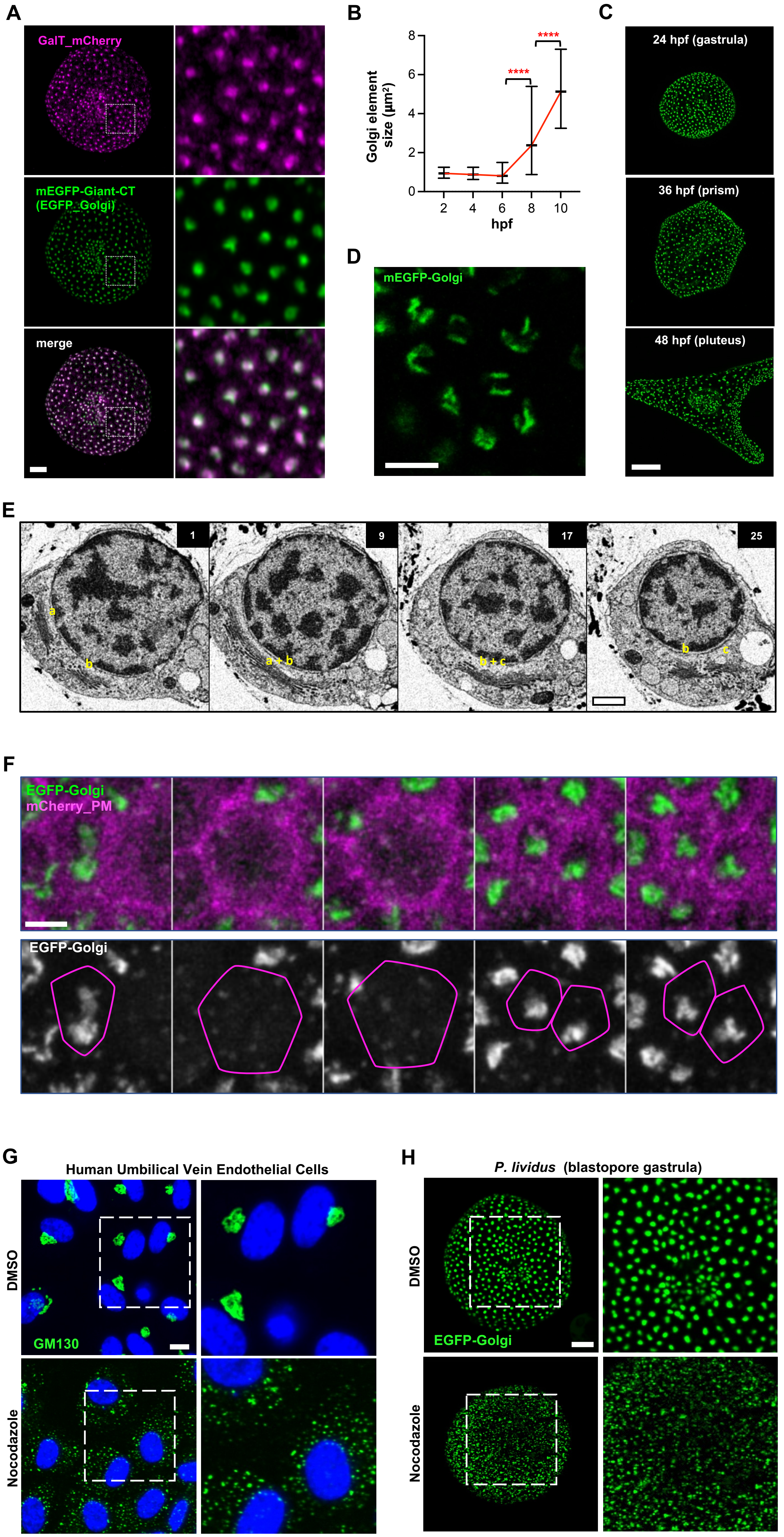

A
GalT_mCherry
mEGFP-Giant-CT
(EGFP_Golgi)
merge
B
Golgi element size (µm²)
8
6
4
2
0
****
****
2
4
6
8
10
hpf
C
24 hpf (gastrula)
36 hpf (prism)
48 hpf (pluteus)
D
mEGFP-Golgi
E
1
9
17
25
a
b
a + b
b + c
b
c
F
EGFP-Golgi
mCherry_PM
EGFP-Golgi
G
Human Umbilical Vein Endothelial Cells
DMSO
GM130
Nocodazole
H
P. lividus (blastopore gastrula)
DMSO
EGFP-Golgi
Nocodazole
